# Water-Only Look-Locker Inversion recovery (WOLLI) T_1_ mapping

**DOI:** 10.1101/2022.01.28.478209

**Authors:** Liam D. Garrison, Christina Levick, Michael Pavlides, Thomas Marjot, Ferenc Mozes, Leanne Hodson, Stefan Neubauer, Matthew D. Robson, Christopher T. Rodgers

## Abstract

**Purpose:** Modified Look-Locker Inversion recovery (MOLLI) and Shortened MOLLI (ShMOLLI) T_1_ values can deviate substantially from the water T_1_ in voxels containing large amounts of water and fat. We introduce Water-Only Look-Locker Inversion recovery (WOLLI) to map water T_1_ in tissue containing fat.

**Theory:** WOLLI comprises adiabatic water-selective inversion and several balanced steady-state free precession (bSSFP) readouts, followed immediately by adiabatic fat-selective inversion and bSSFP readout(s). Data are fitted pixel-wise and an adapted Deichmann-Haase saturation-correction gives the water T_1_.

**Methods:** We compared two WOLLI protocols (WOLLI-w7f1 and WOLLI-w7f2f1), ShMOLLI and single-voxel spectroscopy at 3T in: simulations, fat/water phantoms, and 12 subjects’ livers.

**Results:** In simulations with 0-40% fat fraction, T_1_ varied by: <0.5% (WOLLI-w7f1), <12% (WOLLI-w7f2f1), and >100% (MOLLI). In phantoms, the accuracy of the methods (i.e. worst-case T_1w_ difference vs spectroscopy) was: WOLLI-w7f1 (best, maximum difference of 10.2%), WOLLI-w7f2f1 (max. 10.5%), and ShMOLLI (max. 81.5%). In the liver, the root-mean-squared deviations vs spectroscopy were: 374ms (WOLLI-w7f1), 645ms (WOLLI-w7f2f1), and 700ms (ShMOLLI).

**Conclusion:** Compared to ShMOLLI, WOLLI is less influenced by fat, but more by B_0_-inhomogeneity. WOLLI was a clear improvement in phantoms but less so in vivo. WOLLI may be suited to scan organs with high fat content such as the liver.

## Introduction

T_1_-mapping is a powerful approach for in vivo tissue characterization (1–7). In the heart, T_1_ changes can reveal edema (8), myocardial infarction (9) and fibrosis (10). In the liver, T_1_-mapping can detect the onset of fibrosis or cirrhosis without the risks of biopsy (11,12). Pulse sequences such as Modified Look-Locker Inversion Recovery (MOLLI) (13), its variant Shortened Modified Look-Locker Inversion Recovery (ShMOLLI) (14), and Saturation recovery single-shot acquisition (SASHA) (15) are becoming established for T_1_ mapping in vivo.

In disease, changes in tissue microstructure cause corresponding changes in the T_1_ of water (referred to as the “water T_1_” or T_1w_ below). For example, after myocardial infarction, increased interstitial volume due to edema or rupture of cell membranes causes elevated water T_1_ (16). However, in organs such as the liver, disease progression is often also associated with fat deposition. In severe disease, non-adipose tissue fat fractions can be as high as 40%, often distributed in intracellular lipid droplets (17–21). Popular T_1_ mapping methods do not discriminate between water and fat when inverting (or saturating) the magnetization, or during readout. For voxels containing a mixture of fat and water, this means that the observed T_1_ doesn’t only reflect the water T_1_ in MOLLI, ShMOLLI or SASHA. Kellman *et al.* showed that the water T_1_ can be *overestimated* with even small fat fractions for MOLLI and SASHA in the heart (22); Mozes *et al.* showed this for MOLLI in the liver (23); and Larmour *et al.* showed this in skeletal muscle (24). Differences in fat fraction between subjects, or changes over time, could confuse the interpretation of T_1_ maps and limit their diagnostic and prognostic value.

Ultimately, we hypothesize that a measurement of the fat and a T_1w_ value provides more information on tissue microstructure than a MOLLI T_1_ value that incorporates influences from both fat and water. And hence we hypothesize that T_1w_ values will be more robust markers of disease, especially in studies that involve weight loss (or gain).

New approaches are needed to selectively measure the water T_1_ in tissue even in the presence of fat. Pagano *et al.* introduced the “IDEAL-T_1_” sequence at 1.5T (25), which extends SASHA by adding readouts with multiple echo times (TEs). These are processed by the “iterative decomposition of water and fat with echo asymmetry and least squares estimation” (IDEAL) algorithm to give separate fat and water saturation recovery curves in each pixel, which are then fitted to give a fat T_1_ map and a water T_1_ map from a 2 breath-hold scan. Larmour *et al.* proposed applying MOLLI or SASHA at 1.5T at several frequency offsets each experiencing different fat-induced shifts in the observed T_1_, and which can be used as input in a look-up table to determine fat fraction and the water T_1_ (24). Nezerfat *et al.* adapted the “slice-interleaved T_1_” (STONE) sequence at 1.5T (26) by using an amplitude-modulated pulse to selectively invert water (27–29). This “water-selective slice-interleaved T_1_” (STONE-WS) sequence was described in two abstracts (27,28).

The aim of this study is to investigate whether the fat sensitivity of MOLLI T_1_ measurements can be reduced using frequency-selective inversion of water in a new “Water-Only Look-Locker Inversion recovery” (WOLLI) pulse sequence. We select an appropriate frequency-selective inversion pulse; we show that frequency-selective inversion alone is not sufficient to obtain water T_1w_ via the Deichmann-Haase saturation-correction formula (30) used in MOLLI post-processing (31); and we show how to overcomes this challenge either by gathering a fat-suppressed image, or by fitting the fat T_1_*. We validate WOLLI at 3T using simulations, and by comparing the fat-sensitivity and performance of WOLLI and ShMOLLI relative to spectroscopic reference data in a 19-vial fat/water phantom, and in 12 volunteers’ livers.

## Theory

### Frequency-selective inversion pulse

The MOLLI T_1_-mapping pulse sequence, shown in Figure 1a, comprises one or more “inversion epochs”. Each epoch contains an adiabatic inversion pulse, and a train of single-shot balanced steady-state free precession (bSSFP) readouts that sample the recovering magnetization (14,32). MOLLI is normally run during a breath-hold with electrocardiogram (ECG) gating. The vendor’s implementation of MOLLI on our “3T TIM Trio” scanner (Siemens, Erlangen, Germany) uses a hyperbolic secant (HS) adiabatic inversion pulse with duration T_p_ = 10.16 ms, time-bandwidth product R = 5.439, and truncation = 6.5% (33). Figure 2a shows that this pulse inverts magnetization over γΔB_0_ ≈ ±500Hz.

**Figure 1.**
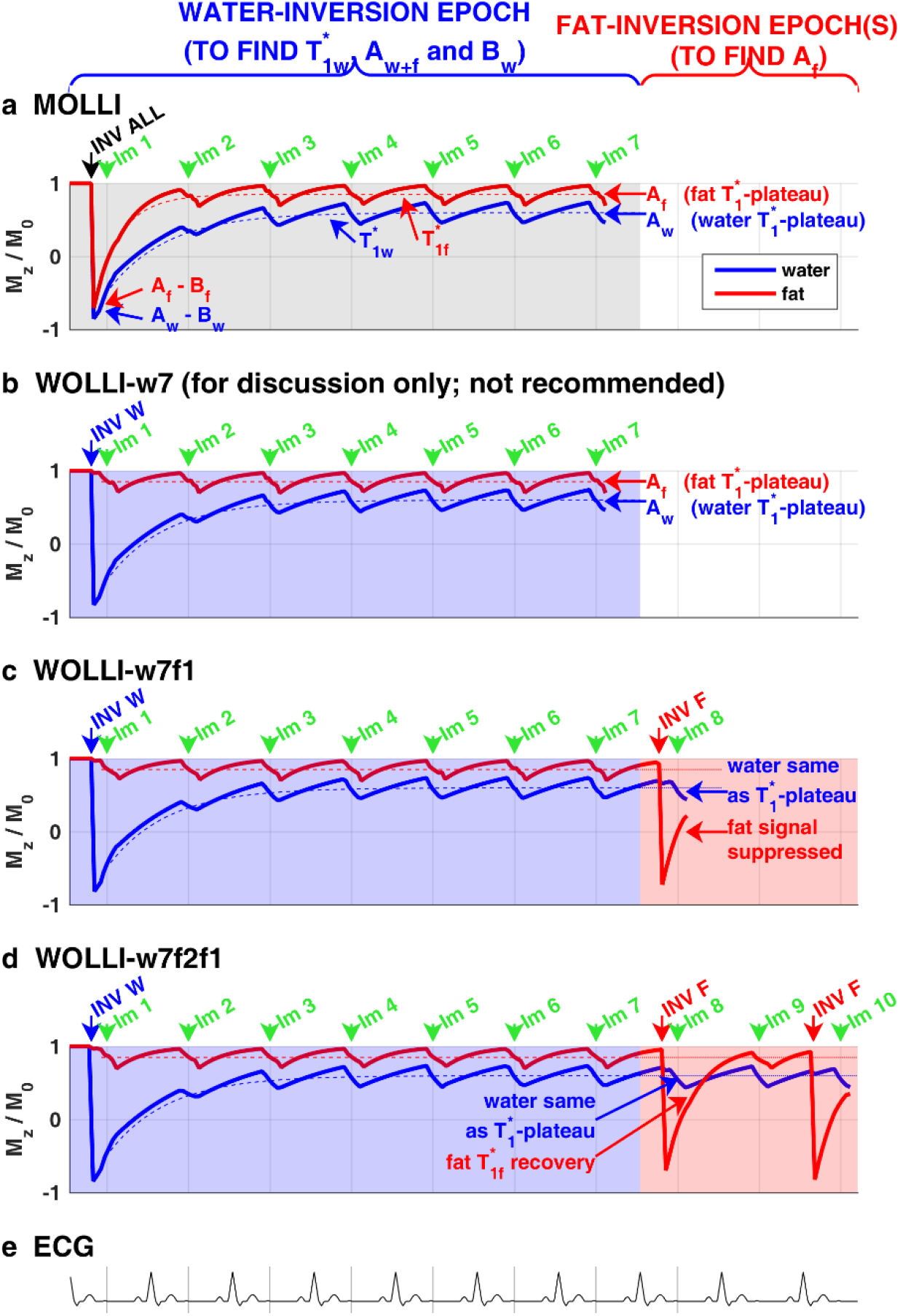
Pulse sequence diagrams comparing longitudinal magnetization of water (blue) and fat (red) for: (a) MOLLI with hyperbolic secant (HS) inversion and a 7-heartbeat non-selective-inversion epoch (shaded grey); (b) WOLLI-w7 which differs from (a) by use of a water-selective inversion pulse; (c) WOLLI-w7f1 with hypergeometric (HG) inversion, a 7-heartbeat water-inversion epoch (shaded in blue), and a fat-suppressed image (shaded in pink); (d) WOLLI-w7f2f1 with hypergeometric (HG) inversion, a 7 heartbeat water-inversion epoch (shaded in blue), and two fat-inversion epochs (shaded in pink) at 228 ms, 228+RR ms and 100 ms TI. (e) Schematic electrocardiogram. Dashed lines in (a-d) illustrate the 3-parameter model for water and fat pools from Eq. 5. Other parameters: 340 ms trigger delay, TI 100+n x RR ms, single-shot bSSFP readout. 11cm width

**Figure 2.**
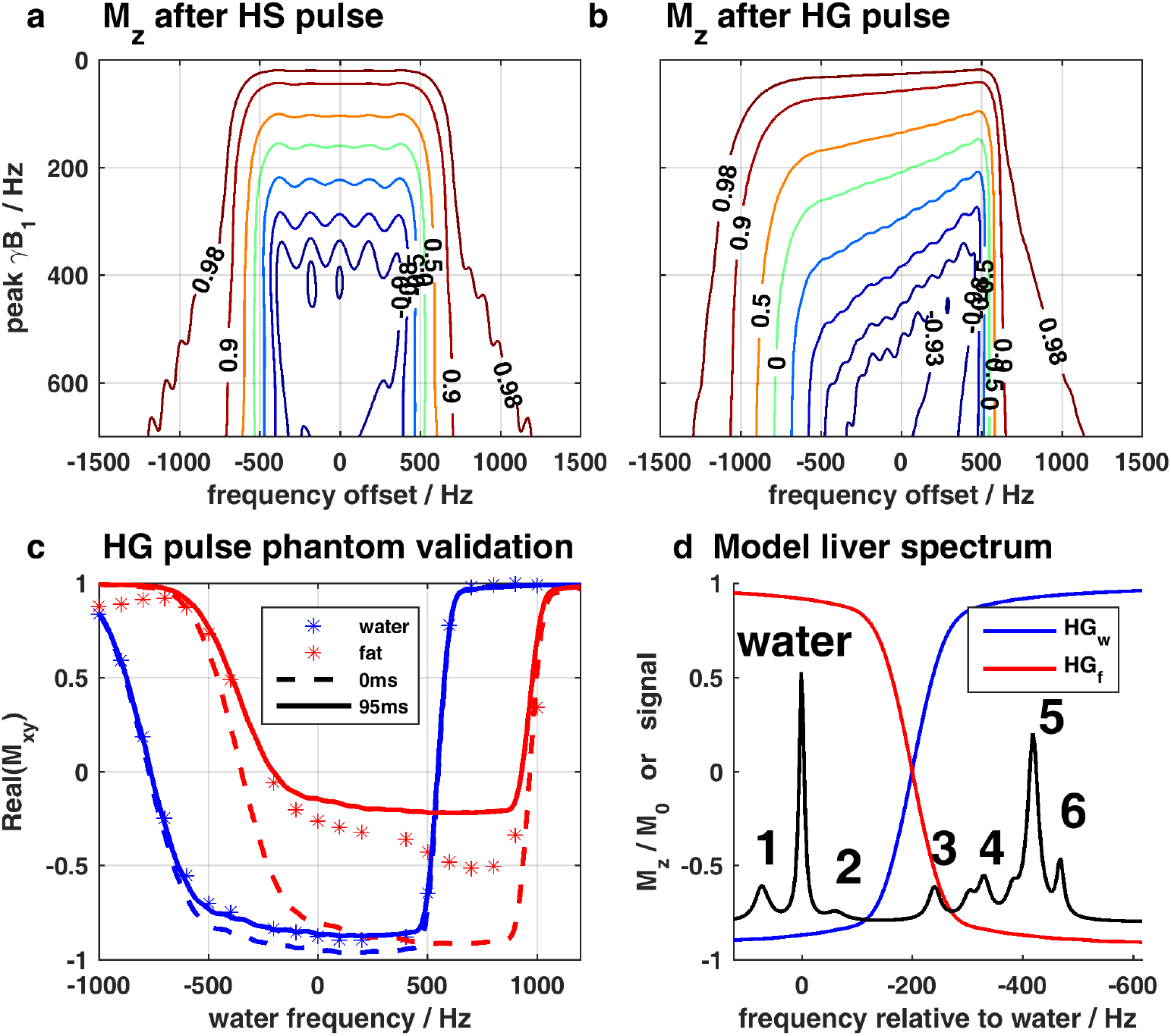
Inversion pulses. (a-b) Contour plots of longitudinal magnetization after the hyperbolic secant inversion used by MOLLI (a), or the hypergeometric (HG) inversion used by WOLLI (b). (c) Phantom validation of HG inversion performance in water/agarose (blue) and fat (red) vials. Note that the fat signal experiences significant T_1_ relaxation between inversion and readout as expected. (d) Simulated fat spectrum as described by Hamilton *et al* (34) overlaid with water-selective (HG_w_, blue) and fat-selective (HG_f_, red) pulse profiles at γB_1_ = 500 Hz, showing the selectivity of these pulses. 1.5-column width

Figure 2d shows the peaks typically seen in a liver spectrum. The principal fat resonance (methylene, or -CH_2-_) occurs at 1.30 ppm, and the water resonance is at 4.70 ppm (34). These peaks differ in frequency by *v*_w_ – *v*_f_ = 420 Hz on the 3T Trio scanner. So the HS pulse inverts fat as well as water.

We sought an adiabatic pulse with similar duration, specific absorption rate (SAR) and adiabatic onset to the MOLLI HS pulse, and which inverts over the range of frequencies *v*_water_ ± γΔB_0_ Hz covering the water resonance, but which has negligible effects over the range of frequencies *v*_fat_ ± γΔB_0_ Hz covering the fat resonance. To quantify this frequency-selectivity, we define the inversion efficiency 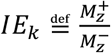, where 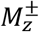 is the magnetization of species *k* immediately after/before the pulse. We sought a pulse with *IE_water_* ≈ –1.0 and *IE_fat_* ≈ +1.0 for fat. In other words, we sought an inversion pulse with a sharp transition-width, a broad passband, and an achievable adiabatic onset. Such a pulse would perform frequency-selective inversion, and would tolerate the B_0_- and B_1_-inhomogeneity typical in the body at 3T (35).

Rosenfeld *et al.* (36) introduced hypergeometric (HG) adiabatic pulses, defined parametrically by:

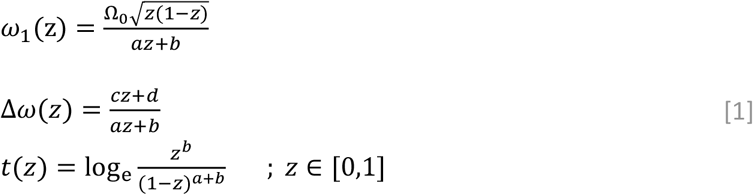

where Ω_0_, *a, b, c* and *d* are shape parameters, *ω*_1_ is the RF amplitude, Δ*ω* is the RF frequency offset, *t* is time, and the parameter *z* sweeps from 0 (at *t* = −∞) to 1 (at *t* = +∞). HG pulses include HS pulses as a special case, but in general HG pulses are *asymmetric* in the time domain and have an asymmetric frequency response. The introduction of asymmetry allows a trade-off between the sharpness of the transition widths on the two sides of the pulse. This allows us to narrow the transition width between fat and water. The broadening of the transition width on the far-side of the pulse will not matter if the passband is wide enough. HG pulses (36) have been used for chemical shift-selective inversion recovery (CSS-IR) fat-water separation (37) and for joint fat and water suppression in spectroscopy (38), but not yet for T_1_-mapping.

### Non-selective inversion recovery

MOLLI typically uses single-shot bSSFP readouts (32). The signal in each voxel is complex-valued, and depends on the off-resonance frequency, flip-angle, echo time, and the T_1_ and T_2_ of the tissue in that voxel (39). In a voxel containing a mixture of water and fat, with a proton-density fat fraction *F_f_*, the complex bSSFP signals sum to give the total voxel signal

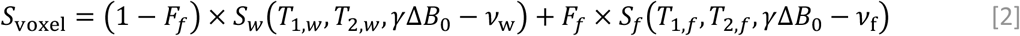

where the subscript “w” denotes water, and “f” denotes fat, and *S_w/f_* are the pure water/fat bSSFP signals, which depends on several tissue parameters.

MOLLI analysis (32) fits the signals *S*_TI_ in each voxel to a 3-parameter model:

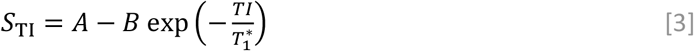

where TI is the time from inversion to the centre of k-space for each image, and the fitted parameters are A (T_1_*-recovery plateau signal), B (A minus the signal immediately after inversion), and 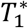 (effective relaxation time). The Deichmann-Haase formula (30) attempts to correct 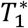 for readout-induced saturation to give:

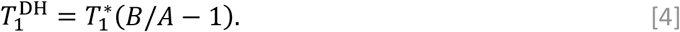

Note that with a typical 2.4/1.2ms TR/TE in MOLLI, *S_w_* and *S_f_* are almost exactly out of phase at 3T (22). This is illustrated in Figure 3a-c by the blue and red markers. Even small amounts of fat can cause fitted T_1_* values to be *greater* than the water T_1_. “Correction” with Eq. 4 then leads to an even higher over-estimation for 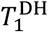.

**Figure 3.**
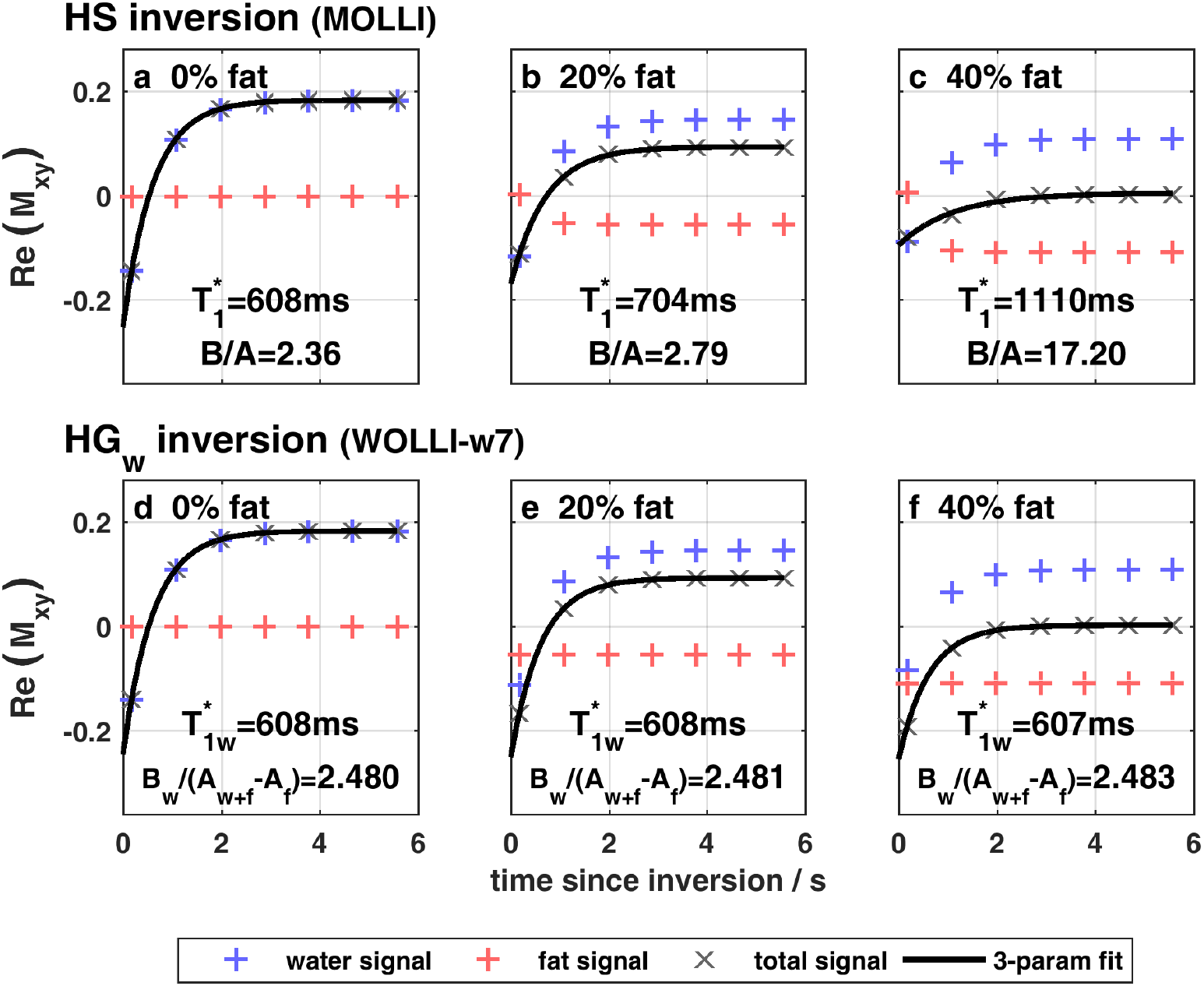
Simulated inversion-recovery curves at three different fat fractions for (a) MOLLI and (b) WOLLI-w7 (i.e. the first seven heartbeats of the WOLLI methods tested later). The total signal (grey crosses) is the weighted sum of the water (blue +) and fat (red +) components. A 3-parameter fit of the total signal (black) is shown with its associated T_1_* and B/A values for Deichmann-Haase correction. These vary with fat fraction for MOLLI, but not for WOLLI. Parameters: 1000ms *T*_1*w*_ 312ms *T*_1*f*_, 35° bSFFP readout, 2.51/1.05ms TR/TE. 1.5-column width

### Frequency-selective inversion recovery

Extending Eq. 3 to treat fat and water separately, the *k*^th^ pool’s contribution to the signal is

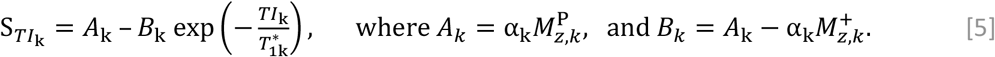

*TI*_k_ is now the time since the *most recent interruption* of the train of readouts for the *k^th^* pool. Note that in a given readout, *TI*_k_ may be different for different pools. E.g. a water-selective inversion pulse resets only *TI*_w_ not *TI*_f_. Other parameters are: the 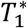-recovery plateau signal from the k^th^ pool *A_k_* (proportional to the longitudinal magnetization 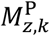); *B_k_* which is analogous to *B* in Eq. 3; the longitudinal magnetization just after the preceding interruption to the train of readouts 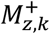; and the apparent relaxation time 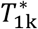. The meaning of these parameters is illustrated in Figure 1.

Let us now consider a system with water and fat magnetization at equilibrium *M*_0,*k*_ that is subjected to a water-selective inversion pulse and a train of readouts. Water is inverted just before the readout train so *IE_w_* ≈ −1; but fat remains at equilibrium so *IE_f_* ≈ +1. Substituting these into Eq. 5 gives: 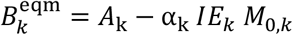. The Deichmann-Haase formula (Eq. 4) can be extended for the k^th^ pool (30,40):

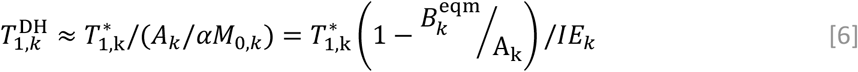

Note that this formula is valid only when the magnetization is initially at equilibrium (as here).

At first glance, it would seem that using a water-selective inversion pulse in a MOLLI-like sequence (e.g. WOLLI-w7 in Fig. 1b), and applying Eq. 6 would give maps of the water *T*_1*w*_. However, careful inspection of Eq. 6 and the blue-shaded region in Figure 1b shows that this is not the case. This is because Deichmann-Haase correction in Eq. 6 requires knowledge of *A_w_,* but the observed *T*_1_*-plateau signal 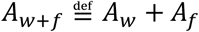 comes from both water that has recovered (*A_w_*, in blue) and fat that was never inverted (*A_f_*, in red).

### Inversion-recovery from the *T*_1_*-plateau

Figure 4 illustrates one approach to separate *A_w_* and *A_f_* so that Eq. 6 can be used to compute *T*_1*w*_. The key idea is to perform inversion-recovery in the middle of a train of readouts. A hypothetical readout at TI = 0ms after an inversion pulse targeted on the k^th^ pool will then give a signal −*A_k_* (blue curve in Fig. 4) instead of the signal 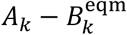 if the k^th^ pool was initially at equilibrium (grey curve). In other words, starting and ending the inversion-recovery at the T_1_*-plateau state eliminates one parameter from Eq. 5, i.e. 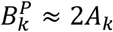.

**Figure 4:**
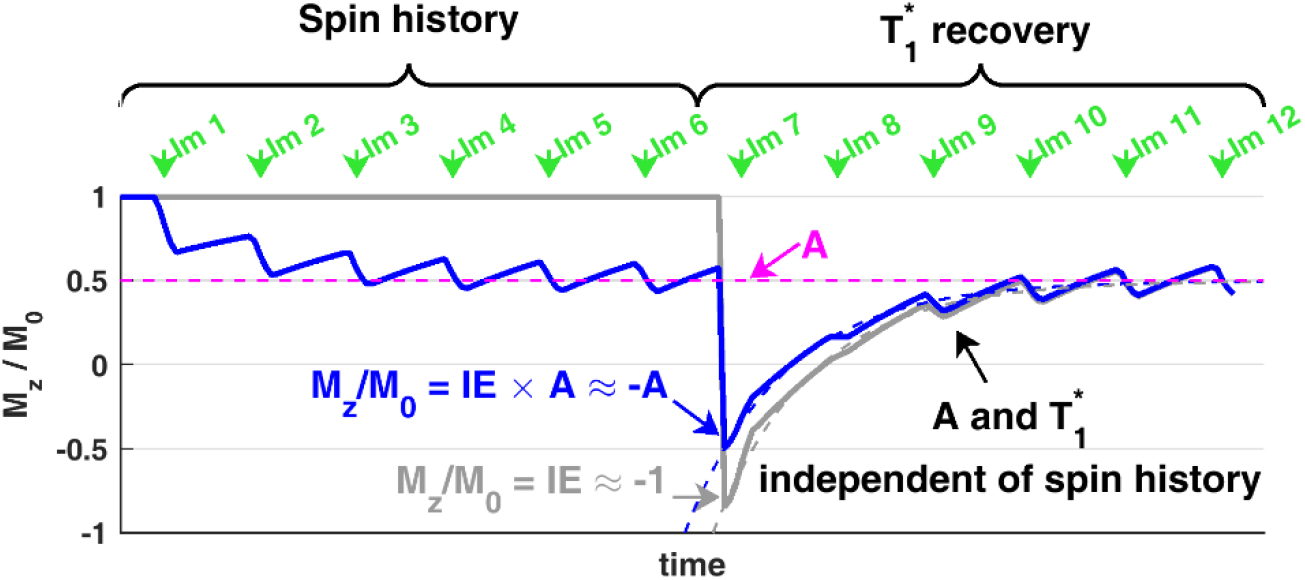
Effect of spin history on the MOLLI 3-parameter model: s = A - B exp(-TI/T_1_*). Grey: inversion follows a 6hb recovery period. Blue: 6 readouts to reach T_1_*-plateau, then inversion, then 6 readouts. Note how inversion from the T_1_*-plateau means that *B* = *A* – *M_z_/M*_0_ ≈ 2*A*. Parameters: 900ms RR interval, 20° bSSFP flip angle, 2000ms *T*_1*w*_ and 17.5ms *T*_2*w*_. 11cm (<1.5-column) width

### WOLLI sequence

The WOLLI sequence (Figs 1c and 1d) applies these two concepts of frequency-selective inversion-recovery and inversion-recovery (for fat) starting from the T_1_*-plateau. We describe the particular protocol by a notation adapted from that used for MOLLI. E.g. WOLLI-w7f2f1 (Fig. 1d) means a WOLLI sequence with water-selective inversion, 7 readouts, fat-selective inversion, 2 readouts, fat-selective inversion and 1 readout. These readouts continue at regular intervals so that the water signal remains at the T_1_*-plateau level during fat inversion-recovery, and 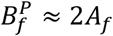. Further epochs could be written as e.g. “WOLLI-w7f2f1(3)w5”, which would add a 3-heartbeat recovery interval, a second water-selective inversion, and five more readouts.

### WOLLI post-processing: “fat T_1_ fitting” algorithm

Consider a WOLLI-w7f2f1 scan as shown in Figure 1d. The fat magnetization remains approximately at its T_1_*-plateau level throughout the water-selective inversion epoch (images 1-7 here) because in liver the fat *T*_l,f_ ≈ 312 ± 48 ms (41) which is much shorter than typical water *T*_1*w*_. Therefore, WOLLI analysis can be performed as follows.

#### Step 1

Use images 1-7 to fit *A*_*f*+*w*_, 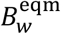, and 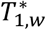 in

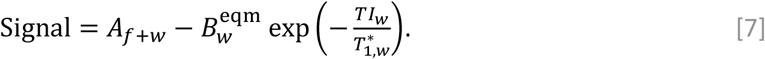

N.B. computing the water *T*_1_ using Eq. 6 requires *A_w_* but this step fits 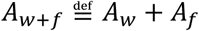 + *A_f_* instead.

#### Step 2

Since 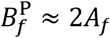, use images 8-10 and the parameters from step 1 to fit 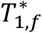 and *A_f_* in

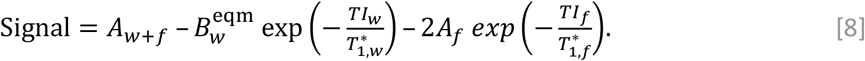

#### Step 3

Regularize *A_f_* in pixels with low fat fraction, where Eq. 8 is underdetermined, either:

a. By updating 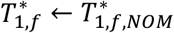 if images 8-10 have a SD < 1% maximum, or if the Jacobian element for 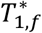 is < 10^-3^ in step 2; or
b. By weighting according to the Cramér-Rao Bound of the fit variance 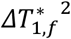

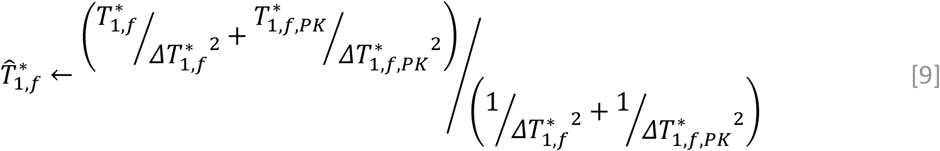

where 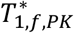 and 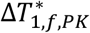 are prior knowledge constants based on Bloch simulations.

#### Step 4

Update *A_f_* by linear least squares in Eq. 8 with all other parameters fixed.

#### Step 5

Compute *A_w_* = *A*_*f*+*w*_ – *A_f_* and hence 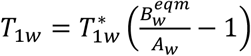 using Eq. 6.

Non-linear fitting is performed using the “lsqcurvefit” implementation of the trust-region reflective algorithm (42,43) in Matlab (The Mathworks Inc, Natick, MA, USA). This algorithm can easily be adapted to other WOLLI protocols.

### WOLLI post-processing: “fat suppression” algorithm

Alternatively, if variation in 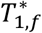 is negligible, *A_w_* can be obtained from a fat-suppressed image. E.g. in WOLLI-w7f1 (Fig. 1c), a fat-selective inversion pulse is timed to null fat in the 8^th^ image while leaving the water signal at the T_1_*-plateau. Substituting 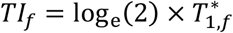 into Eq. 8 and rearranging gives an alternative fitting algorithm.

#### Step 1

*As above.*

#### Step 2

Compute *A*_w_ using 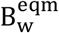 and 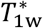 from Step 1 and the formula

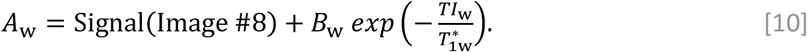

#### Step 3

Compute 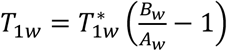 using Eq. 6.

## Methods

### Simulations

Simulations were performed in Matlab using Hargreaves’ Bloch Equation Simulator (from http://www-mrsrl.stanford.edu/~brian/blochsim/) for two non-exchanging pools: water (*v_w_* = 0 Hz) and fat (*v*_f_ = −420 Hz). This corresponds to the field strength of 2.89 T on our “3T Trio” scanner.

Hypergeometric (HG) pulses were optimized for selective inversion starting from parameters that replicated the symmetric hyperbolic secant (HS) inversion pulse in ShMOLLI (14), and using literature liver T_1_ and T_2_ values (41). We simulated a water-selective “HG_w_” pulse and a fat-selective “HG_f_” pulse, obtained by conjugating the HG_w_ pulse and shifting its central frequency. We sought an 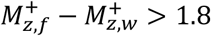 over at least a 200Hz range of γΔB_0_ (to cover the readout bSSFP passband), and an adiabatic onset γB_1_ ≤ 700 Hz. Figure 2b shows a simulated frequency profile of the optimized HG pulse, which had Ω_0_=4.7, a=1, b=0.3, c=-6, d=1.5, and 2% truncation. Figure 2c shows phantom validation of this profile. Figure 2d shows the predicted effect of M_z_ in vivo with HG_w_ shifted by +350Hz, and HG_f_ conjugated and shifted by −700Hz to place the transition bands for both pulses between water and fat resonances.

We then simulated 7-heartbeat MOLLI, WOLLI-w7f1, and WOLLI-w7f2f1. These differ only in the choice of inversion pulse and in the fat-selective inversion epoch(s) in WOLLI. We used the bSSFP shaped pulses and timings from the MOLLI “WIP561” package (Siemens), including the appropriate HS pulse (for MOLLI) or our optimized HG pulses (for WOLLI). Parameters were: 900 ms RR interval, T_1,w_ = 1000 ms, T_1,f_ = 312 ms, T_2,w_ = 40 ms, and T_2,f_ = 60 ms (41), and bSFFP readout: 35° flip angle, TR/TE = 2.51 ms/1.05 ms, 5 ramp-up pulses, 24 full pulses to centre of k-space, 66 full pulses in total, 1 ramp-down pulse.

### Imaging protocol

Scans were acquired using a 3T Trio scanner. B_0_ shimming used the gradient recalled echo shimming “WIP452B” package (Siemens, Erlangen, Germany).

ShMOLLI T_1_ mapping used the MOLLI “WIP561” package (Siemens, Erlangen, Germany), with online reconstruction by the method of Piechnik *et al* (14,44), and the liver T_1_ mapping protocol of Banerjee *et al* (11). We used: a ShMOLLI 5(1)1(1)1 protocol, with HS inversion (T_p_ = 10.16 ms, R = 5.439, truncation = 6.5%), TIs of 100ms + n × RR, 180ms and 260ms for the three inversion epochs, and bSSFP readout with 35° flip angle, TR/TE = 2.51ms/1.05ms, 5 ramp-up pulses, 24 full pulses to center of k-space, 66 full pulses in total, 1 ramp-down pulse, and with 1.9×1.9×8.0 mm^3^ voxel size, 360 x 270 x 8mm^3^ field of view, 192×144 acquisition matrix, 2x generalized auto-calibrating partially parallel acquisition (GRAPPA) acceleration (45), 6/8 partial Fourier and adaptive coil combination (46).

The WOLLI-w7f1 protocol (Fig. 1c) had: (a) an HG_w_ pulse in heart-beat 1, (b) seven readouts with 100ms + n × RR TI in heart-beats 1-7, (c) an HG_f_ pulse in heart-beat 8, and (d) one readout with 228ms TI in heart-beat 8.

The WOLLI-w7f2f1 protocol (Fig. 1d) had: (a-c) as above, (d) two readouts with 100ms and 100ms + RR TI in heart-beats 8-9, (e) an HG_f_ pulse in heart-beat 10, and (f) one readout with 218 ms TI in heart-beat 10.

WOLLI T_1w_-maps were processed offline in MATLAB.

Reference water T_1_ values were recorded with a custom inversion recovery stimulated echo acquisition mode (STEAM-IR) single voxel spectroscopy (SVS) sequence, with: 90° flip angle, 2 kHz bandwidth, 10 ms TE, 7 ms mixing time (TM), no water suppression, 1024 vector size, 1 preparation scan, and whitened singular value decomposition (WSVD) coil combination (47,48). Spectra were analyzed in Matlab, by fitting the complex signals with the advanced method for accurate, robust, and efficient spectral fitting (AMARES) (49). Then the complex water peak amplitudes were phased and fitted to a 3-parameter model [Signal = *A* – *B* exp(−*TI/T*_1,*w*_)].

Proton-density fat fraction (PDFF) was quantified using a custom sequence (50), following an established liver protocol (11). Spectra were acquired with water suppression (STEAM, 2s TR, 3 averages, 10ms TE, 7ms TM) and without water suppression (4s TR, 1 average). PDFF was then determined using AMARES in MATLAB.

### Phantom study

Mixtures of water (18MΩ purity), fat (peanut oil, TESCO, UK), agarose (2% bw), NaCl, NiCl_2_, sodium dodecyl sulphate, sodium benzoate (all from Sigma-Aldrich, UK) were emulsified in 30mL vials in an Ultrasonic Cleaner (UD200SH-6LQ, Eumax, Hong Kong). Nineteen vials were placed in a polypropylene cylinder (275mm 0, 130mm h, Sealfresh Ltd, UK) filled with 50 mM CuSO_4(aq)_ solution. 18 vials had T_1w_ of approximately 700 ms, 1100 ms and 1400 ms and proton-density fat fractions of 0%, 5%, 10%, 20%, 30%, 40%. The 19^th^ vial contained only fat.

Imaging was performed as described above in a coronal slice through the centre of the phantom. STEAM-IR was run with 30s TR, 3 averages, and 14×14×40mm^3^ voxels centered over each vial in turn.

### In-vivo liver study

This study was approved by our institution’s Research Department and the National Research Ethics Service. All volunteers gave informed written consent. Imaging with ShMOLLI, WOLLI-w7f1 and WOLLI-w7f2f1 was performed as described above in a transverse slice through the livers of 12 subjects (8 males, 4 females; height: 1.67 ± 0.17 m; weight: 87 ± 18 kg; body mass index (BMI): 32 ± 9; age: 46 ± 14 years; PDFF: 9 ± 7 %). Ten subjects were patients at high risk of non-alcoholic fatty liver disease (NAFLD) – being overweight (BMI > 25) and resistant to insulin (having a fasting plasma insulin concentration in the top 10% of the Oxford Biobank range (51)). Reference T_1_ values were acquired with STEAM-IR, and fat fraction with STEAM-PDFF acquired from 20×20×20mm^3^ voxels intersecting the imaging slice, in the lateral part of the right lobe of the liver, away from major vessels, bile ducts and the edge of the liver. STEAM-IR used a 10s TR, 1 average, and 50, 1218, 2385, 3553, 4720 ms TIs. STEAM-PDFF was as above.

## Results

### Simulations

Figure 3 compares 7hb MOLLI and WOLLI-w7, i.e. with HS and HG_w_ inversion respectively. Water T_1_* values vary by 83% for fat fractions 0–40% with MOLLI, but only by 0.2% with WOLLI. The ratio B/A varies from 2.36 to 17.20 with MOLLI, and so the “correction” for partial saturation in Eq. 4 introduces large errors in T_1w_ measurement in the presence of fat, whereas *B_w_*/(*A*_w+f_ – *A_f_*) varies by 0.1%, so WOLLI avoids these errors. Note, that B_eqm_/A_w+f_ = 2.33, 3.67 and 80.74 for the three WOLLI plots. This is very different to the value of ≈ 2 that is expected for Deichmann-Haase correction, and shows clearly the need to distinguish between A_w_ and A_w+f_ when correcting for partial saturation.

Figure 5 shows the errors in fitted water T_1w_ values for MOLLI, WOLLI-w7f1 and WOLLI-w7f2f1. At 0% Ff, all three methods show an approximately linear dependence of fitted vs true T_1w_ from 160-2000 ms. MOLLI underestimates T_1w_ by 21%, WOLLI-w7f1 by 12%, and WOLLI-w7f2f1 by 15%. This underestimation is expected, being largely due to imperfect inversion (14). In the presence of up to 30% fat, WOLLI-w7f1 and WOLLI-w7f2f1 retain their modest linear underestimation of T_1w_, whereas MOLLI shows large, non-linear changes in T_1_.

**Figure 5:**
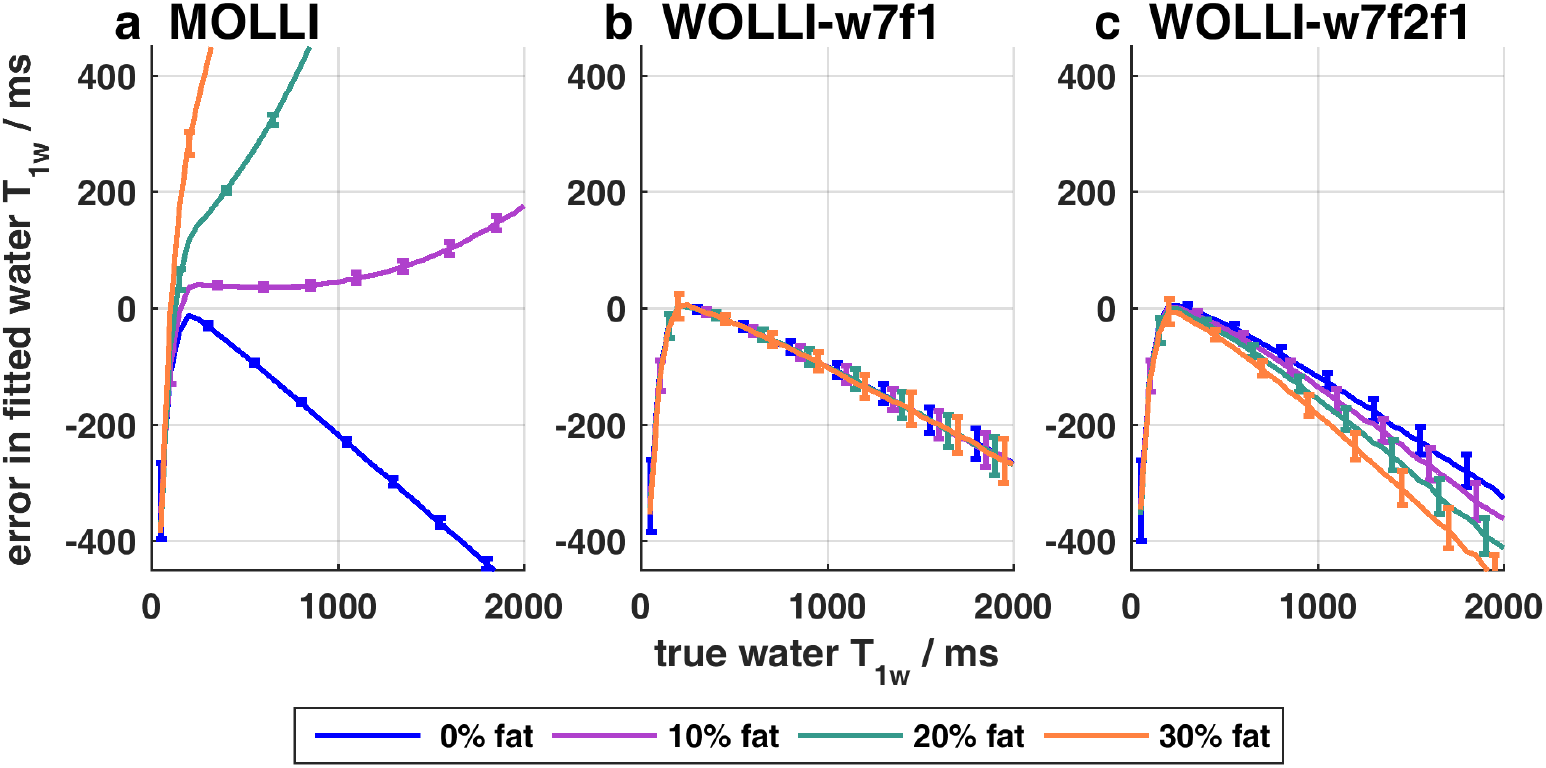
T_1_ accuracy and precision of MOLLI (a) and WOLLI (b-c). Mean ± st. dev. of fitted T_1_ (or T_1w_) are plotted for a range of input water T_1w_ values (0–2000ms) and fat fractions (0%, 10%, 20%, 30%). The simulation ran 200 repetitions at an SNR = 250. SNR was defined as the magnitude of the simulated T_1_*-plateau signal level for 100% fat, divided by the standard deviation of the noise added to the real and imaginary parts of the simulated pixel values. 1.5-column width

Figure 6 shows that WOLLI T_1_ values are consistent up to 30% fat fraction for off-resonance frequencies from −100 to +100 Hz. This is similar to the frequency range of MOLLI’s bSSFP readout. The WOLLI off-resonance response is asymmetric due to the HG inversion pulses.

**Figure 6.**
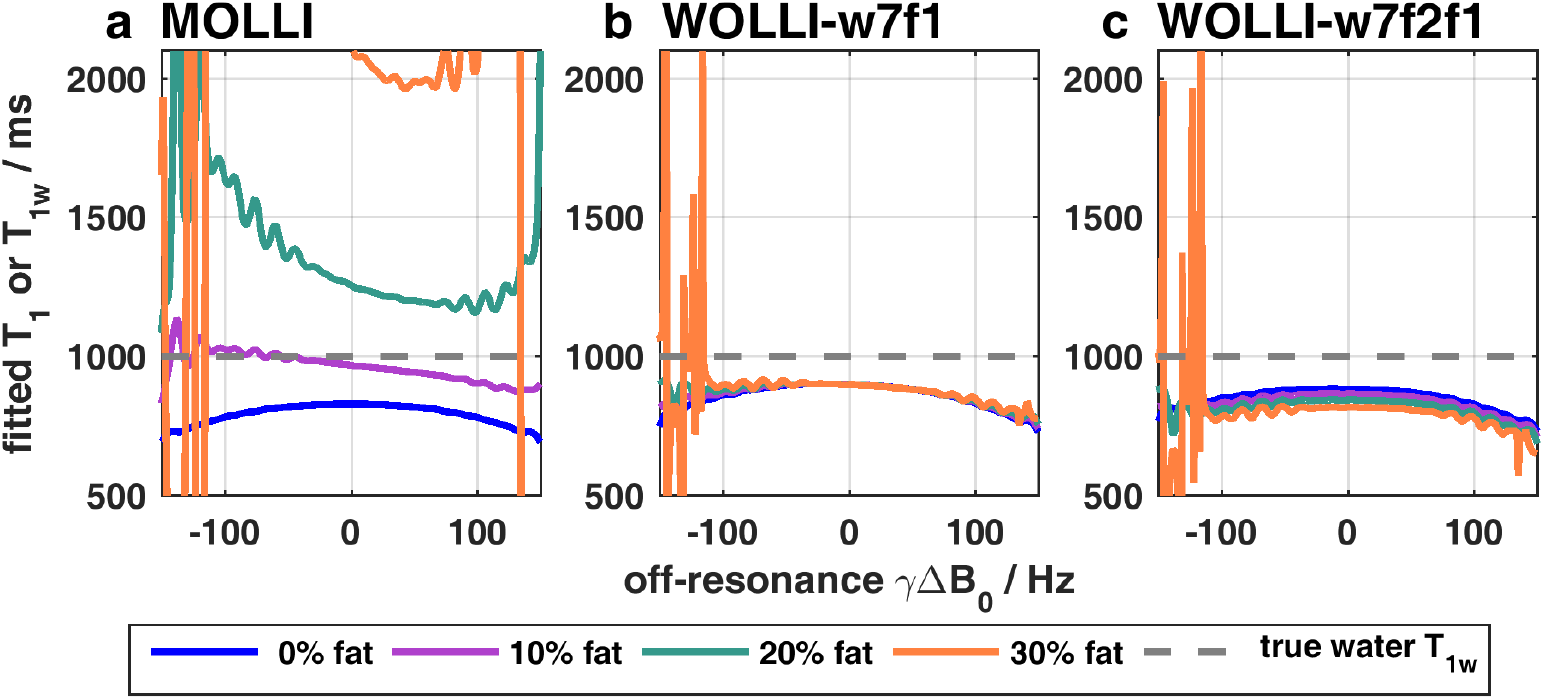
Off-resonance performance of (a) MOLLI and (b-c) WOLLI for 0, 10, 20, 30% fat fractions. 1.5-column width.

Figure 7 shows Monte Carlo simulations to test the accuracy (mean) and precision (standard deviation, SD) of MOLLI and WOLLI T_1_ values. The mean T_1_ varies negligibly with SNR from 100 to 300. For a signal-to-noise ratio (SNR) of 250, which is typical for human liver scans, and with *Ff* = 0%, the T_1_ SDs of WOLLI were approximately twice that for MOLLI (MOLLI: 5.7, WOLLI-w7f1 : 12.6, and WOLLI-w7f2f1: 13.9). When *F_f_* = 20%, the T_1_ SDs became similar (MOLLI: 12.3, WOLLI-w7f1 : 14.9, WOLLI-w7f2f1: 17.5). When *F_f_* = 30%, the T_1_ SD of MOLLI is much greater than WOLLI.

**Figure 7.**
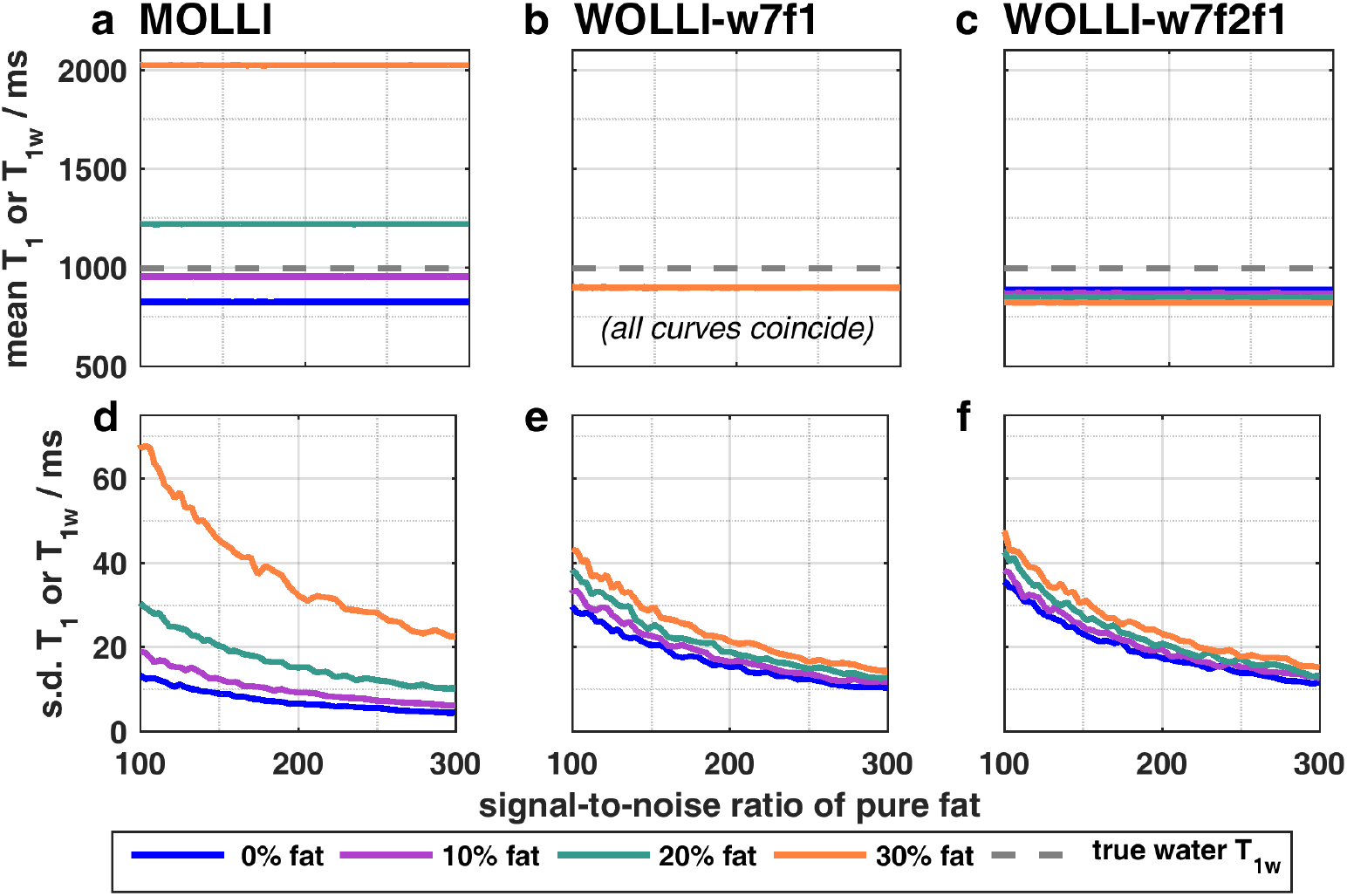
T_1_ accuracy (mean, a-c) and precision (st. dev., d-f) of fitted T_1_ (or T_1w_) as a function of signal-to-noise ratio for MOLLI (a,d) and WOLLI (b,c,e,f). The simulation ran 1000 repetitions at each of 60 signal-to-noise ratios (100–300) and 4 fat fractions (0–30%). SNR was defined as the magnitude of the simulated T_1_*-plateau signal level for 100% fat, divided by the standard deviation of the noise added to the real and imaginary parts of the simulated pixel values. 1.5-column width.

### Phantom study

Figure 2c shows the HG pulse profile measured in the vials containing pure water and pure fat. For frequency offsets from −1000 to +1000 Hz, the maximum error for water was 0.78% of the equilibrium signal. For fat, it was 30%, due to differences in T_1_ and T_2_ of peanut oil in the phantom compared to the human liver literature values used for simulations.

Figure 8 compares T_1_ values from ShMOLLI, WOLLI-w7f1, WOLLI-w7f2f1 and the STEAM-IR reference method in the phantom. The STEAM-IR reference T_1_ values in the 3 groups of vials with equal aqueous NiCl_2_ concentration were 658 ± 7ms, 1136 ± 22ms, and 1369 ± 7ms (mean ± inter-vial SD). WOLLI-w7f1 T_1_ values differed from the STEAM-IR reference values by a minimum of 0.4% to a maximum of 10.2% across the vials; WOLLI-w7f2f1 T_1_ values differed by 0.8% to 10.5%; and ShMOLLI T_1_ values differed by 0.6% to a maximum of 81.5%.

**Figure 8.**
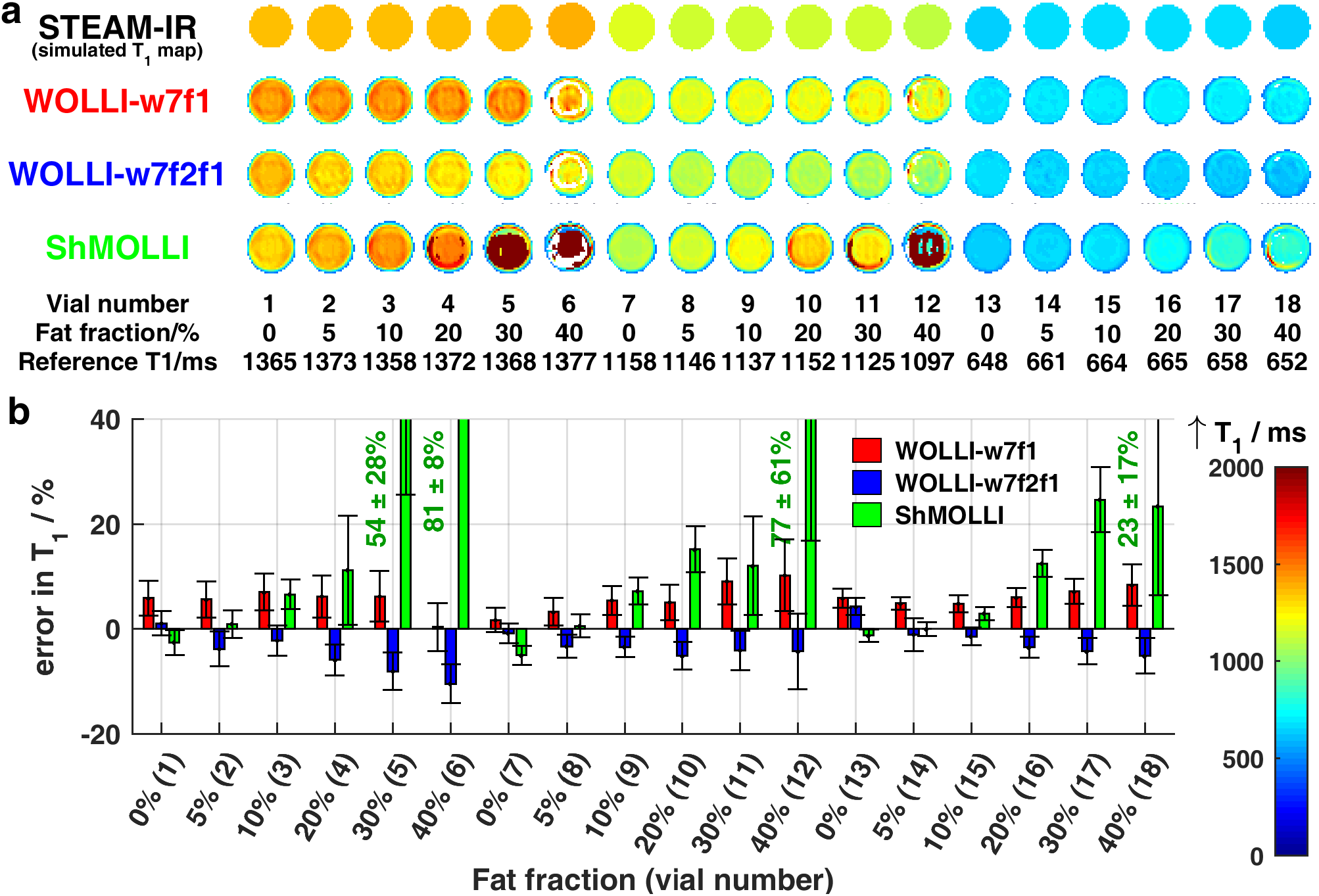
Multi-compartment phantom results. (a) T_1_ maps for each method and each vial are shown in a montage. STEAM-IR reference water T_1_ values are also plotted in the same color scale. Fat fractions are indicated for each vial. (b) Percentage error from STEAM-IR reference T_1_ for each of the methods, and normalized standard deviation of T_1_ values. These mean and standard deviation values were taken from a circular region of interest in the center of each tube. The 19^th^ (pure fat) vial is not shown because it contained too little water to measure the water T_1_. 2-column width

### In-vivo liver study

Figure 9 summarizes the results from scans in the livers of 12 subjects with PDFF in the range 4 - 26%, by plotting measured T_1w_ values against STEAM-IR reference T_1w_ values. WOLLI-w7f1 had the least deviation from the line of identity, with a root mean squared deviation (RMSD) of 374 ms, compared to 645 ms (WOLLI-w7f2f1), and 700 ms (ShMOLLI). Specifically, WOLLI-w7f1 T_1_ values differed from the STEAM-IR reference values by a minimum of 6.3% to a maximum of 20.3%; WOLLI-w7f2f1 by 3.0% to −35.0%; and ShMOLLI by 2.1% to 28.8%. WOLLI T_1_ maps from a typical subject (Fig. 9e-g) show the expected abdominal anatomy.

**Figure 9.**
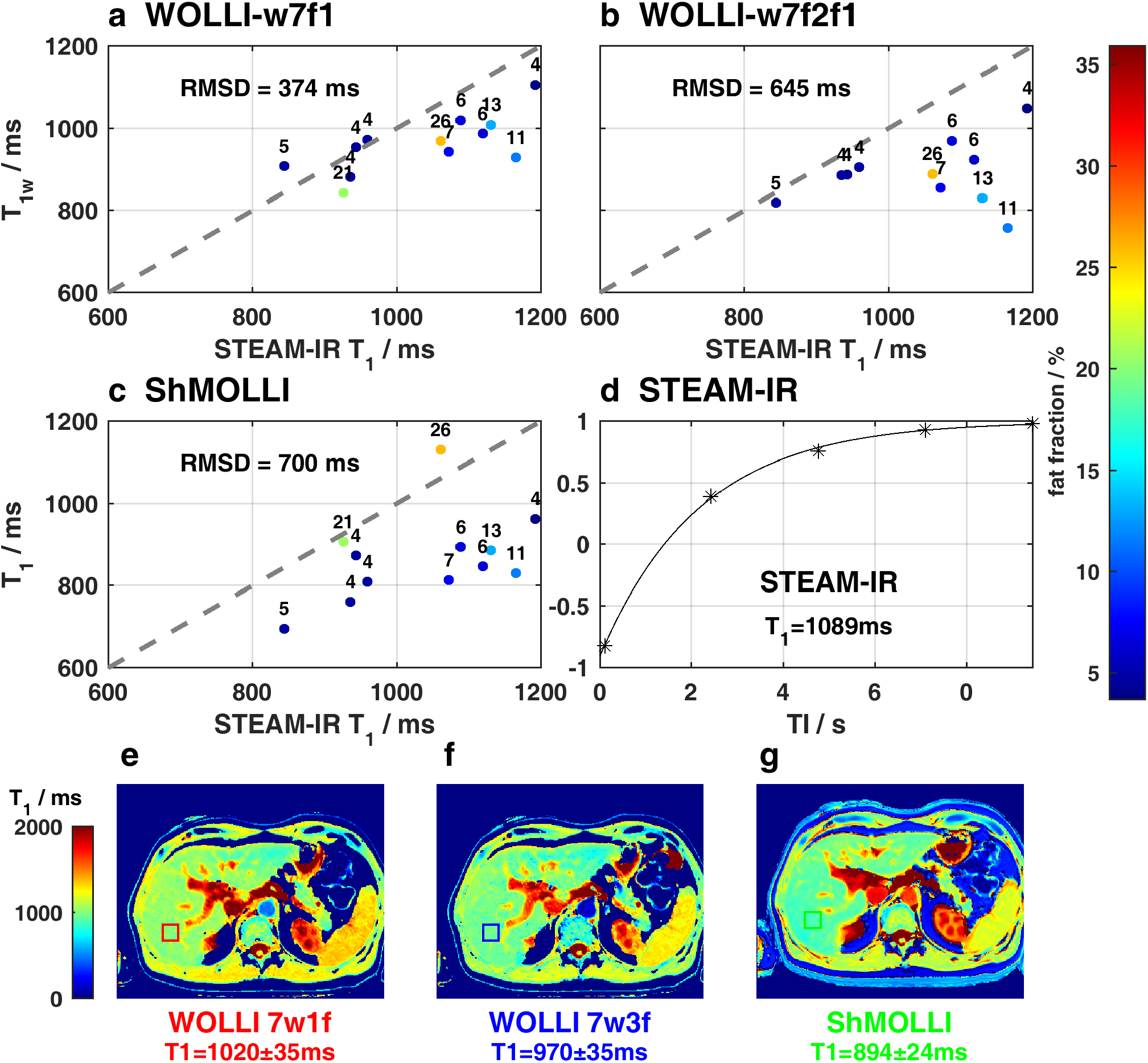
Performance in human liver. (a-c) Scatter plots of measured T_1_ vs STEAM-IR reference T_1_. Subjects are labelled with their associated percentage fat fraction, and colored accordingly. The root mean squared deviation (RMSD) is from the line of identity (grey). (d) STEAM-IR recovery curve and (e-g) T_1_ maps from a 67-year-old male subject. The STEAM-IR voxel is marked on the T_1_ maps as a colored square.

## Discussion

### Simulations

HG inversion pulses can be used at 3T to selectively invert water or fat with similar pulse duration and peak B1 requirements, as shown in Figure 2. Substituting the HS global inversion pulse in MOLLI for a water-selective HG_w_ pulse enables robust measurement of T_1w_* in the presence of fat, as shown in Figure 3. However, to obtain T_1w_ it is necessary to separate the contribution to the T_1_*-plateau signal level of water (A_w_) from that of non-inverted fat (A_f_). We did this either by recording a fat-suppressed image (e.g. WOLLI-w7f1) or by adding fat-selective inversion-recovery epoch(s) (e.g. WOLLI-w7f2f1). WOLLI gives acceptable accuracy in T_1w_ measurements for a range of water T_1_ values and fat fractions (Figure 5), and frequency offsets (Figure 6). WOLLI has greatly increased accuracy (error in T_1w_ mean) in the presence of fat than MOLLI, but at a cost of lower precision (T_1w_ SD) than MOLLI at low fat fractions (Figure 7).

### Phantom study

In phantoms, the accuracy of the methods (assessed by worst-case T_1w_ differences) were in the order: WOLLI-w7f1 (best, maximum difference of 10.2%), WOLLI-w7f2f1 (max. 10.5%), and ShMOLLI (max. 81.5%). WOLLI has significantly reduced the effect of fat on the measured T_1_, but in its present form WOLLI has not achieved complete independence of measured T_1_ from the fat fraction.

### In-vivo liver study

The in-vivo T_1_ accuracy plotted in Figure 9 shows the same ordering, with WOLLI-w7f1 best, then WOLLI-w7f2f1, and finally ShMOLLI. However, the performance of WOLLI in vivo is less impressive than the simulations and phantom study suggest it could have been.

### Limitations

In this study, we modelled fat using only the methylene peak, which constitutes 70% of the total fat signal in the liver (34). More accurate simulations could have incorporated all 6 fat resonances. Nevertheless, the validation of WOLLI in the phantom (with all fat peaks present), and in vivo supports our choice to use a two pool model, which is easier to interpret. Note that to minimize the impact of using a 2-pool model, the HG_w_ pulse was placed to minimally perturb most fat resonances, and the HG_f_ pulse was placed to invert most fat resonances as shown in Figure 2d.

Our WOLLI fitting algorithm was designed to be numerically robust and simple enough to implement online for future studies. In step 1 of the WOLLI fitting algorithms, we assume that the fat signal is constant throughout the water inversion epoch (images 1-7). However, this is not entirely true, particularly for the first image, as seen in Figure 3. However, since the slight change in amplitude in image 1 does not fit the model in Eq. 8, the effect on the fitted parameters is negligible. In principle, this approximation could be avoided by fitting the 6 parameters: 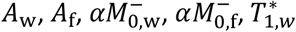, and 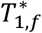,> but we found that 6-parameter Levenberg-Marquardt fitting did not converge robustly. A more robust reconstruction approach, e.g. dictionary-matching magnetic resonance fingerprinting (52) based on Bloch simulations (53), might improve the performance of WOLLI for future studies.

Another limitation relates to variability of the fat T_1f_, T_2f_ and hence the fat inversion efficiency and effective relaxation time constant 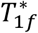. Figure 10 compares the performance of MOLLI with three WOLLI protocols as a function of fat fraction, and for the expected T_1f_ value and at ±15%. All the WOLLI protocols give the water T_1w_ more accurately than MOLLI for PDFF >5%.

**Figure 10.**
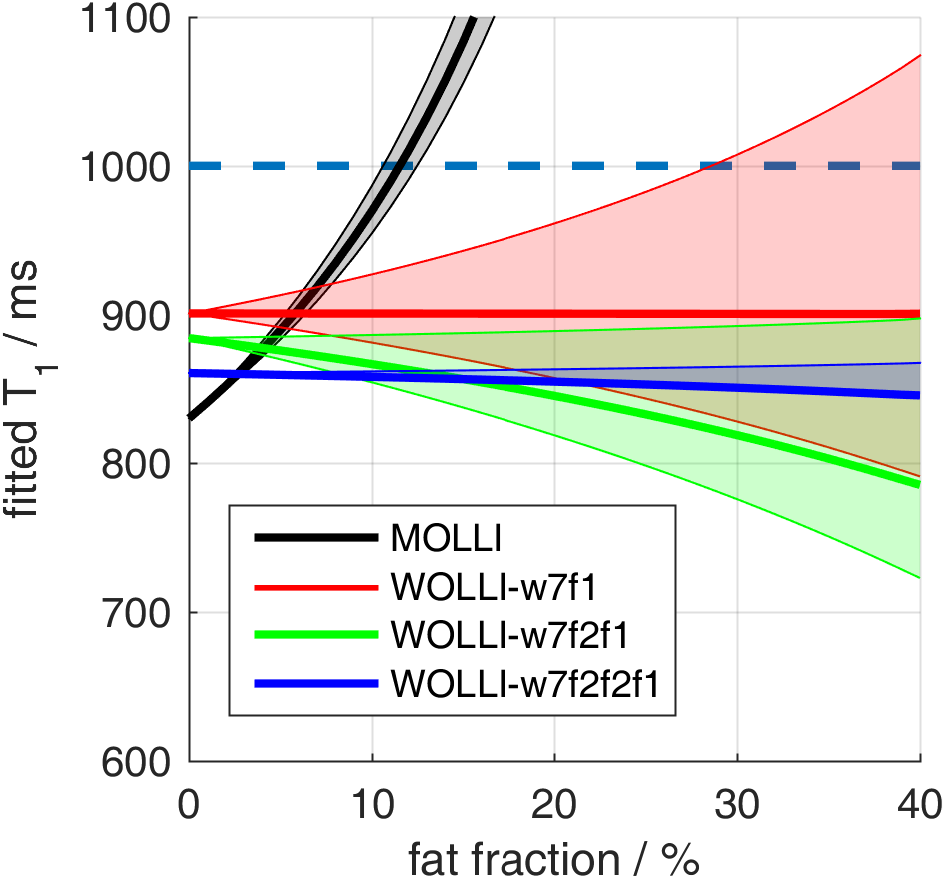
Effect of increasing fat fraction and fat T_1f_ on the different T_1_ mapping methods. Simulations used the liver fat T_1f_ from Hamilton *et al.* (34) (thick line) and T_1f_ values ±15% of this (thin lines / shaded area). MOLLI fits extremely high T_1_ values for 15-35% fat fractions, and fitting does not even converge for 35-60% fat fractions due to the cancellation of water and fat signals. 1-column width

If the fat T_1f_ is known (thick lines), then WOLLI-w7f1 T_1w_ values depend very little on fat fraction (varying by <0.5% for fat fractions from 0 to 40%), compared to WOLLI-w7f2f1 (varying by <12%) and MOLLI (varying by >> 100%). But if the fat T_1_ deviates from its expected value (thin lines / shaded area) then WOLLI-w7f1 will be less desirable because it has the strongest dependence of T_1w_ on T_1f_ of all the methods. It might be better in such circumstances to use an alternative fat suppression method before the final image, such as a train of lower amplitude HG_f_ pulses, analogous to ref. (54), that could suppress fat robustly over a wider range of T_1f_ than simple fat-inversion-recovery.

We have focused on two representative sampling schemes (WOLLI-w7f1 and WOLLI-w7f2f1) to illustrate the concept of water-selective inversion-recovery in WOLLI. A great many sampling schemes are possible. The accuracy and precision of T_1w_ depends on the detailed interplay between the choice of sampling scheme and the approximations in the WOLLI fitting algorithms. For example, if it is acceptable to lengthen the breath-hold to 12 heart-beats then WOLLI-w7f2f2f1 (shown in blue in Fig. 10) has 5 data points to determine fat T_1f_, and hence is predicted to be significantly more robust to fat T_1f_ variation. Or if a study requires post-contrast T_1w_ mapping, we would recommend using an optimized WOLLI sampling strategy based on further simulations.

## Conclusions

Swapping to a water-selective inversion pulse in MOLLI is not enough to obtain water T_1w_ maps. This is because the correction for readout-induced saturation (from T_1_* to T_1_) must disregard the background signal from fat. Saturation-correction can be achieved using an additional fat-suppressed image (in WOLLI-w7f1) or fat-selective inversion-recovery epoch (in WOLLI-w7f2f1).

In simulations and phantoms, WOLLI T_1w_ values were less sensitive to fat than MOLLI T_1_ values, particularly for fat fractions >20%. In human livers, WOLLI reduced the root-mean-squared deviation of T_1(w)_ from 700ms (ShMOLLI) to 374ms (WOLLI-w7f1) or 645ms (WOLLI-w7f2f1) vs spectroscopy.

WOLLI is slightly more sensitive to B_0_-inhomogeneity than MOLLI, and WOLLI T_1_ maps have higher T_1_ SD in tissue with low fat fractions (<10%) than MOLLI maps.

WOLLI provides an attractive alternative approach for T_1_-mapping in tissue that is suspected to contain ≥5% fat, such as diseased livers. Further studies are now required to test whether WOLLI T_1w_ will be a clinically useful tissue biomarker.

## Acknowledgements

CTR is funded by a Sir Henry Dale Fellowship from the Wellcome Trust and the Royal Society [098436/Z/12/B and 098436/Z/12/Z]. CTR acknowledges support from the Cambridge NIHR Biomedical Research Centre [BRC-1215-20014]. We thank William Clarke for providing Bloch simulation tools and Aaron Hess for helpful discussion. This work is the subject of US Patent US10386429B2 with priority date 22 Apr 2016.

## Notes

### Competing Interest Statement

The work described in this manuscript is the subject of US patent US10386429B2.

## References

1. Moon J, Abdel-Gadir A, Treibel T. Myocardial T1 mapping: where are we now and where are we going? Res. Reports Clin. Cardiol. 2014:339. doi: 10.2147/RRCC.S50891.

2. Jellis CL, Kwon DH. Myocardial T1 mapping: modalities and clinical applications. Cardiovasc. Diagn. Ther. 2014;4:126–37. doi: 10.3978/j.issn.2223-3652.2013.09.03.

3. Kramer CM, Chandrashekhar Y, Narula J. The Tissue Issue T1 Mapping and the Myocardium. JACC Cardiovasc. Imaging 2016;9:88–90. doi: 10.1016/j.jcmg.2015.11.003.

4. Kramer CM, Chandrashekhar Y, Narula J. T1 mapping by CMR in cardiomyopathy: A noninvasive myocardial biopsy? JACC Cardiovasc. Imaging 2013;6:532–534. doi: 10.1016/j.jcmg.2013.02.002.

5. Bellm S, Basha TA, Shah R V, Murthy VL, Liew C, Tang M, Ngo LH, Manning WJ, Nezafat R. Reproducibility of Myocardial T 1 and T 2 Relaxation Time Measurement Using Slice-Interleaved T 1 and T 2 Mapping Sequences. 2016:1–9. doi: 10.1002/jmri.25255.

6. Moon JC, Messroghli DR, Kellman P, et al. Myocardial T1 mapping and extracellular volume quantification: a Society for Cardiovascular Magnetic Resonance (SCMR) and CMR Working Group of the European Society of Cardiology consensus statement. J. Cardiovasc. Magn. Reson. 2013;15. doi: Artn 92Doi 10.1186/1532-429x-15-92.

7. Schelbert EB, Messroghli DR. State of the Art: Clinical Applications of Cardiac T1 Mapping 1. 2016;278.

8. Ferreira VM, Piechnik SK, Dall’Armellina E, Karamitsos TD, Francis JM, Choudhury RP, Friedrich MG, Robson MD, Neubauer S. Non-contrast T1-mapping detects acute myocardial edema with high diagnostic accuracy: a comparison to T2-weighted cardiovascular magnetic resonance. J. Cardiovasc. Magn. Reson. 2012;14:42. doi: 10.1186/1532-429X-14-42.

9. Dall’Armellina E, Piechnik SK, Ferreira VM, et al. Cardiovascular magnetic resonance by non contrast T1-mapping allows assessment of severity of injury in acute myocardial infarction. J. Cardiovasc. Magn. Reson. 2012;14:15. doi: 10.1186/1532-429X-14-15.

10. Bull S, White SK, Piechnik SK, et al. Human non-contrast T1 values and correlation with histology in diffuse fibrosis. Heart 2013;99:932–937. doi: 10.1136/heartjnl-2012-303052.

11. Banerjee R, Pavlides M, Tunnicliffe EM, et al. Multiparametric magnetic resonance for the non-invasive diagnosis of liver disease. J. Hepatol. 2014;60:69–77. doi: 10.1016/j.jhep.2013.09.002.

12. Neuberger J. Guidelines on the use of Liver Biopsy in Clinical Practice. BSG Guidel. Gastroenterol. 2004:1–15. doi: 10.1136/gut.45.2008.iv1.

13. Messroghli DR, Radjenovic A, Kozerke S, Higgins DM, Sivananthan MU, Ridgway JP. Modified Look-Locker inversion recovery (MOLLI) for high-resolutionT1 mapping of the heart. Magn. Reson. Med. 2004;52:141–146. doi: 10.1002/mrm.20110.

14. Piechnik SK, Ferreira VM, Dall’Armellina E, Cochlin LE, Greiser A, Neubauer S, Robson MD. Shortened Modified Look-Locker Inversion recovery (ShMOLLI) for clinical myocardial T1-mapping at 1.5 and 3 T within a 9 heartbeat breathhold. J. Cardiovasc. Magn. Reson. 2010;12:1–11. doi: 10.1186/1532-429x-12-69.

15. Chow K, Flewitt J a., Green JD, Pagano JJ, Friedrich MG, Thompson RB. Saturation recovery single-shot acquisition (SASHA) for myocardial *T*_1_ mapping. Magn. Reson. Med. 2014;71:2082–2095. doi: 10.1002/mrm.24878.

16. Dall’Armellina E, Piechnik S, Ferreira V, et al. Cardiovascular magnetic resonance by non contrast T1-mapping allows assessment of severity of injury in acute myocardial infarction. J. Cardiovasc. Magn. Reson. [Internet] 2012;14:15.

17. Takahashi Y, Fukusato T. Histopathology of nonalcoholic fatty liver disease/nonalcoholic steatohepatitis. World J. Gastroenterol. 2014;20:15539–15548. doi: 10.3748/wjg.v20.i42.15539.

18. Brunt EM, Tiniakos DG. Histopathology of nonalcoholic fatty liver disease. World J. Gastroenterol. 2010;16:5286–5296. doi: 10.3748/wjg.v16.i42.5286.

19. McCrae J, Klotz O. The Distribution of Fat in the Liver. J Exp Med [Internet] 1910;12:746–754.

20. Delikatny EJ, Chawla S, Leung DJ, Poptani H. MR-visible lipids and the tumor microenvironment. NMR Biomed. 2011;24:592–611. doi: 10.1002/nbm.1661.

21. Martin S, Parton RG. a Dynamic Organelle. Mol. Cell [Internet] 2006;7:373–378. doi: 10.1038/nrm1912.

22. Kellman P, Bandettini WP, Mancini C, Hammer-Hansen S, Hansen MS, Arai AE. Characterization of myocardial T1-mapping bias caused by intramyocardial fat in inversion recovery and saturation recovery techniques. J. Cardiovasc. Magn. Reson. 2015;17:33. doi: 10.1186/s12968-015-0136-y.

23. Mozes FE, Tunnicliffe EM, Pavlides M, Robson MD. Influence of Fat on Liver T 1 Measurements Using Modified Look – Locker Inversion Recovery (MOLLI) Methods at 3T. 2016:1–7. doi: 10.1002/jmri.25146.

24. Larmour S, Chow K, Kellman P, Thompson RB. Characterization of T 1 bias in skeletal muscle from fat in MOLLI and SASHA pulse sequences: Quantitative fat-fraction imaging with T 1 mapping. Magn. Reson. Med. [Internet] 2016;0:n/a-n/a. doi: 10.1002/mrm.26113.

25. Pagano JJ, Chow K, Yang R, Thompson RB. Fat-water separated myocardial T1 mapping with IDEAL-T1 saturation recovery gradient echo imaging. J. Cardiovasc. Magn. Reson. 2014;16:P65. doi: 10.1186/1532-429X-16-S1-P65.

26. Weingärtner S, Roujol S, Akçakaya M, Basha TA, Nezafat R. Free-Breathing Multislice Native Myocardial T_1_ Mapping Using the Slice-Interleaved T_1_ (STONE) Sequence. Magn. Reson. Med. 2015;124:115–124. doi: 10.1002/mrm.25387.

27. Nezafat M, Roujol S, Jang J, Basha T, Botnar RM. Eliminating the Impact of Myocardial Lipid Content on Myocardial T1 Mapping Using a Spectrally-Selective Inversion Pulse. In: Proc ISMRM. ; 2015. p. 2638.

28. Nezafat M, Roujol S, Jang J, Basha T, Botnar RM. Myocardial T1 mapping with spectrally-selective inversion pulse to reduce the influence of fat. J Cardiovasc Magn Reson 2016;18:18–20. doi: 10.1186/1532-429X-18-S1-P19.

29. Havla L, Basha T, Rayatzadeh H, Shaw JL, Manning WJ, Reeder SB, Kozerke S, Nezafat R. Improved fat water separation with water selective inversion pulse for inversion recovery imaging in cardiac MRI. J. Magn. Reson. Imaging 2013;37:484–490. doi: 10.1002/jmri.23779.

30. Deichmann R, Haase a. Quantification of T1 Values by SNAPSHOT-FLASH NMR Imaging. J. Magn. Reson. 1992;96:608–612. doi: 10.1016/0022-2364(92)90347-A.

31. Messroghli DR, Walters K, Plein S, Sparrow P, Friedrich MG, Ridgway JP, Sivananthan MU. Myocardial T-1 mapping: Application to patients with acute and chronic myocardial infarction. Magn. Reson. Med. 2007;58:34–40. doi: Doi 10.1002/Mrm.21272.

32. Messroghli DR, Radjenovic A, Kozerke S, Higgins DM, Sivananthan MU, Ridgway JP. Modified Look-Locker inversion recovery (MOLLI) for high-resolution T-1 mapping of the heart. Magn. Reson. Med. 2004;52:141–146. doi: Doi 10.1002/Mm.20110.

33. Kellman P, Herzka D a., Hansen MS. Adiabatic inversion pulses for myocardial T1 mapping. Magn. Reson. Med. 2014;71:1428–1434. doi: 10.1002/mrm.24793.

34. Hamilton G, Yokoo T, Bydder M, Cruite I, Schroeder ME, Sirlin CB, Middleton MS. In vivo characterization of the liver fat 1H MR spectrum. NMR Biomed. 2011;24:784–790. doi: 10.1002/nbm.1622.

35. Sung K, Nayak KS. Design and use of tailored hard-pulse trains for uniformed saturation of myocardium at 3 Tesla. Magn. Reson. Med. 2008;60:997–1002. doi: Doi 10.1002/Mrm.21765.

36. Rosenfeld D, Panfil SL, Zur Y. Design of adiabatic pulses for fat-suppression using analytic solutions of the Bloch equation. Magn. Reson. Med. 1997;37:793–801.

37. Laurent WM, Bonny JM, Renou JP. Imaging of water and fat fractions in high-field MRI with multiple slice chemical shift-selective inversion recovery. J. Magn. Reson. Imaging 2000;12:488–496. doi: 10.1002/1522-2586(200009)12:3<488::AID-JMRI15>3.0.CO;2-5.

38. Zhu H, Ouwerkerk R, Barker PB. Dual-band water and lipid suppression for MR spectroscopic imaging at 3 Tesla. Magn. Reson. Med. 2010;63:1486–1492. doi: 10.1002/mrm.22324.

39. Bieri O, Scheffler K. Fundamentals of balanced steady state free precession MRI. J. Magn. Reson. Imaging 2013;38:2–11. doi: 10.1002/jmri.24163.

40. Rodgers CT, Piechnik SK, DelaBarre LJ, Van de Moortele P-F, Snyder CJ, Neubauer S, Robson MD, Vaughan JT. Inversion Recovery at 7 T in the Human Myocardium: Measurement of T-1, Inversion Efficiency and B-1(+). Magn. Reson. Med. 2013;70:1038–1046. doi: 10.1002/mrm.24548.

41. Hamilton G, Middleton MS, Hooker JC, Haufe WM, Forbang NI, Allison M a., Loomba R, Sirlin CB. In vivo breath-hold ^1^ H MRS simultaneous estimation of liver proton density fat fraction, and *T* 1 and *T* 2 of water and fat, with a multi-TR, multi-TE sequence. J. Magn. Reson. Imaging [Internet] 2015:n/a-n/a. doi: 10.1002/jmri.24946.

42. Coleman TF, Li Y. An Interior Trust Region Approach for Nonlinear Minimization Subject to Bounds. SIAM J. Optim. 1996;6:418–445. doi: 10.1137/0806023.

43. Coleman TF, Li Y. On the convergence of interior-reflective Newton methods for nonlinear minimization subject to bounds. Math. Program. 1994;67:189–224. doi: 10.1007/BF01582221.

44. Piechnik SK, Ferreira VM, Lewandowski AJ, et al. Normal variation of magnetic resonance t1 relaxation times in the human population at 1.5t using shMOLLI. J Cardiovasc Magn Reson [Internet] 2013;15:13.

45. Griswold MA, Jakob PM, Heidemann RM, Nittka M, Jellus V, Wang J, Kiefer B, Haase A. Generalized Autocalibrating Partially Parallel Acquisitions (GRAPPA). Magn. Reson. Med. 2002;47:1202–1210. doi: 10.1002/mrm.10171.

46. Walsh DO, Gmitro AF, Marcellin MW. Adaptive reconstruction of phased array MR imagery. Magn. Reson. Med. 2000;43:682–690.

47. Rodgers CT, Robson MD. Receive array magnetic resonance spectroscopy: Whitened singular value decomposition (WSVD) gives optimal bayesian solution. Magn. Reson. Med. 2010;63:881–891. doi: 10.1002/mrm.22230.

48. Rodgers CT, Robson MD. Coil combination for receive array spectroscopy: Are data-driven methods superior to methods using computed field maps? Magn. Reson. Med. [Internet] 2015;0:n/a-n/a. doi: 10.1002/mrm.25618.

49. Vanhamme L, Van Den Boogaart A, Huffel S Van. Improved Method for Accurate and Efficient Quantification of MRS Data with Use of Prior Knowledge. J. Magn. Reson. 1997;129:35–43. doi: 10.1006/jmre.1997.1244.

50. Rial B, Robson MD, Neubauer S, Schneider JE. Rapid quantification of myocardial lipid content in humans using single breath-hold 1H MRS at 3 Tesla. Magn Reson Med [Internet] 2011;66:619–624. doi: 10.1002/mrm.23011.

51. Tan GD, Neville MJ, Liverani E, Humphreys SM, Currie JM, Dennis L, Fielding BA, Karpe F. The in vivo effects of the Pro12Ala PPARg2 polymorphism on adipose tissue NEFA metabolism: The first use of the Oxford Biobank. Diabetologia 2006;49:158–168. doi: 10.1007/s00125-005-0044-z.

52. Ma D, Gulani V, Seiberlich N, Liu K, Sunshine JL, Duerk JL, Griswold MA. Magnetic resonance fingerprinting. (1). Nature [Internet] 2013;495:187–192. doi: 10.1038/nature11971.

53. Shao J, Rapacchi S, Nguyen KL, Hu P. Myocardial T1 mapping at 3.0 tesla using an inversion recovery spoiled gradient echo readout and bloch equation simulation with slice profile correction (BLESSPC) T1 estimation algorithm. J. Magn. Reson. Imaging 2016;43:414–425. doi: 10.1002/jmri.24999.

54. Tao Y, Hess AT, Keith GA, Rodgers CT, Liu A, Francis JM, Neubauer S, Robson MD. Optimized Saturation Pulse Train for Human First-Pass Myocardial Perfusion Imaging at 7T. Magn. Reson. Med. 2015;73:1450–1456. doi: 10.1002/mrm.25262.

